# Maturyoshka: a maturase inside a maturase, and other peculiarities of the novel chloroplast genomes of marine euglenophytes

**DOI:** 10.1101/2021.09.24.461685

**Authors:** Kacper Maciszewski, Nadja Dabbagh, Angelika Preisfeld, Anna Karnkowska

## Abstract

Organellar genomes often carry group II introns, which occasionally encode proteins called maturases that are important for splicing. The number of introns varies substantially among various organellar genomes, and bursts of introns have been observed in multiple eukaryotic lineages, including euglenophytes, with more than 100 introns in their plastid genomes. To examine the evolutionary diversity and history of maturases, an essential gene family among euglenophytes, we searched for their homologs in newly sequenced and published plastid genomes representing all major euglenophytes’ lineages. We found that maturase content in plastid genomes has a patchy distribution, with a maximum of eight of them present in *Eutreptiella eupharyngea*. The most basal lineages of euglenophytes, Eutreptiales, share the highest number of maturases, but the lowest number of introns. We also identified a peculiar convoluted structure of a gene located in an intron, in a gene within an intron, within yet another gene, present in some Eutreptiales. Further investigation of functional domains of identified maturases shown that most of them lost at least one of the functional domains, which implies that the patchy maturase distribution is due to frequent inactivation and eventual loss over time. Finally, we identified the diversified evolutionary origin of analysed maturases, which were acquired along with the green algal plastid or horizontally transferred. These findings indicate that euglenophytes’ plastid maturases have experienced a surprisingly dynamic history due to gains from diversified donors, their retention, and loss.

## 1. Introduction

Group II introns are peculiar, ancient mobile genetic elements, which utilize both autocatalytic RNA and enzymatic peptide components for their own excision, mobility and retrohoming (Lambowitz and Belfort, 2015). Having originated within bacterial genomes, they spread via various means of horizontal gene transfer across the entire tree of life, including mitochondrial and plastid genomes of multiple groups of eukaryotes (Zimmerly and Semper, 2015). During the course of their evolution within the eukaryotic nucleus, group II introns most likely eventually gave rise to more complex spliceosomal introns, although this structurally and functionally major transition still has not been fully explained (Zimmerly and Semper, 2015). It has been hypothesized that the emergence of the spliceosome, together with the formation of the nuclear envelope, are some of the profound evolutionary consequences of the intron invasion of the ancient eukaryotic genome (Lambowitz and Belfort, 2015).

In opposition to the fate of the eukaryotic nucleus-encoded introns, some group II introns followed the path of decreasing complexity, losing parts of their structure and functionality. This is the case group II introns in plant and algal plastid genomes, which are mostly secondarily immobile, and in many cases, only a single intron per plastid genome (ptDNA) has retained its original intron-encoded protein (IEP) – the reverse transcriptase/maturase (Lambowitz and Belfort, 2015; Zimmerly and Semper, 2015). The widespread degeneration of IEPs was possible due to the ability of intron maturases to carry out their duties *in trans*, as it was shown to occur in plant plastids, where all ptDNA-encoded introns are spliced by a single maturase – *matK* (Barthet et al., 2020).

Although the structure, functions and evolutionary history of group II intron and IEPs in bacterial, mitochondrial and plant plastid genomes have been examined in detail for quite some time (Lambowitz and Belfort, 2015; Zimmerly and Semper, 2015), recent years have also brought a substantial number of important studies on their counterparts in non-plant ptDNA (Dabbagh et al., 2017; Janouškovec et al., 2013; Karnkowska et al., 2018). Several algal lineages, bearing both primary (Janouškovec et al., 2013) and secondary plastids (Khan and Archibald, 2008), have been observed to possess group II introns, and among these, the available data points toward their exceptionally convoluted and intriguing evolutionary history in euglenophytes (Dabbagh et al., 2017; Karnkowska et al., 2018; Sheveleva and Hallick, 2004) – descendants of a heterotrophic flagellate host which engulfed a pyramimonadalean green alga (Hrdá et al., 2012). Plastid genomes of contemporary photosynthetic euglenids are not only riddled with highly variable numbers of introns and maturase IEPs, but their functionality is often distinct as well; very few among the plastid group II introns have retained their IEP component, while most have undergone such substantial degeneration that they are currently classified as separate entities – the so-called group III introns (Karnkowska et al., 2018; Zimmerly and Semper, 2015).

Following the acquisition of the plastid, euglenophytes have undergone considerable diversification, forming two main subgroups – the marine order Eutreptiales and the freshwater order Euglenales – and a broad variety of forms, characterized by distinct morphological traits and lifestyles (Kostygov et al., 2021). Marine euglenids, in comparison to the freshwater ones, are a functionally understudied group, most likely due to their substantially lower diversity (constituting only two genera – *Eutreptia* and *Eutreptiella*) and abundance in the environment (Lukešová et al., 2020). Consequently, certain fundamental questions about the biology and evolution of euglenophytes remain unanswered. For instance, the characteristics of the pyramimonadalean ancestor of the euglenid plastid, as well as the identity of its closest extant relative, have not been fully understood (Hrdá et al., 2012; Karnkowska et al., 2018). Moreover, the phylogenetic relationships of Eutreptiales – with Euglenales and each other – are still debated (Karnkowska et al., 2018; Kolisko et al., 2020; Marin et al., 2003).

This work’s primary objective is to investigate the evolutionary history of plastid genomes of marine euglenophytes, with a particular focus on their uncommon traits, such as group II introns and their maturases. Our secondary objective is to utilize the newly-obtained genomic data to unravel the currently uncertain phylogeny of the extant strains classified as members of the genera *Eutreptia* and *Eutreptiella*. With this data, we aim to describe the early history of the euglenophytean secondary plastid, the origins of their fascinating genetic peculiarities and the characteristics of the earliest-branching subdivisions of the photosynthetic euglenids, which may be the key to unraveling the evolutionary paths of Euglenophyta as a whole.

## 2. Materials and Methods

### 2.1. Cultivation and isolation of the genetic material

Culture strains of *Eutreptiella* sp. CCMP389 and *Eutreptiella* sp. CCMP1594 were obtained from the NCMA culture collection (formerly CCMP; National Center for Marine Algae and Microbiota, Bigelow, ME, USA) and maintained in modified L1-Si medium (Guillard and Hargraves, 1993) with artificial seawater Sea-Pure (CaribSea, Inc. Fort Pierce), recommended by the culture collection, under 12/12-hour light/dark cycle at 20-23 °C. After three weeks 50 ml of the culture was centrifuged (2 mins, 4000 rpm). The culture strain *Eutreptiella eupharyngea* K-0027 was obtained from the Norwegian Culture Collection of Algae (NORCCA; Norwegian Institute for Water Research, Oslo, Norway) and cultivated on TL30 medium for marine algae, provided by the culture collection along with the strain. The culture was maintained under 16/8-hour light/dark cycle at 4 °C for one month; afterwards, two 15 ml replicates of the culture were centrifuged (2 mins, 3000 rpm). The culture strain *Eutreptia* sp. CCAC1914B was obtained from the Central Collection of Algal Cultures (CCAC; University of Duisburg-Essen, Germany), and a 10 ml replicate of the culture was centrifuged (2 mins, 3000 rpm).

Cell pellets of *Eutreptiella* sp. CCMP1594, *Etl. eupharyngea* and *Eutreptia* sp. CCAC1914B were subjected to total DNA isolation using DNeasy Blood & Tissue Kit (QIAGEN, USA), following the manufacturer’s protocol, and RNA digestion using RNase A. A subsequent quality control by spectrophotometric measurement of DNA concentration and purity in the samples was performed using an Implen NP80 NanoPhotometer (Implen GmbH, Germany). Total DNA of *Eutreptiella* sp. CCMP389 was isolated with My-budget DNA Mini Kit (Bio-Budget Technologies) following the manufacturer’s protocol. For ensuring sufficient quality and quantity of DNA, a NanoDrop 2000 Spectrophotometer (Thermo Fisher Scientific) was used to measure the concentration and purity of extracted DNA based on A260/A280 and A260/A230 ratios.

### 2.2. High-throughput DNA sequencing and plastid genome assembly

Total DNA samples of *Eutreptiella* sp. CCMP1594, *Etl. eupharyngea* and *Eutreptia* sp. CCAC1914B were handed to an external company (Genomed S.A., Warsaw, Poland) for high-throughput sequencing using Illumina MiSeq platform. The sequencing yielded approximately 6.2 million 250 bp-long paired-end reads for *Eutreptiella* sp. CCMP1594, 3.8 million 300 bp-long paired-end reads for *Etl. eupharyngea*, and 6.7 million 300 bp-long paired-end reads for *Eutreptia* sp. Total DNA sample of *Eutreptiella* sp. CCMP389 was subjected to high-throughput sequencing and assembly by Eurofins Genomics (Ebersberg, Germany), using Illumina HiSeq and the sequencing yielded approximately 4.0 million 150 bp-long paired-end reads. Quality control of the obtained libraries was carried out by the FastQC tool v0.11.6 (Andrews, 2010). Data trimming included filtering of low-quality reads (mean Phred value <15) and removal of the Illumina Universal Adapters, and was carried out using Trimmomatic v0.39 (Bolger et al., 2014).

Genome assembly of the three investigated strains was carried out using SPAdes v.3.10.1 (Nurk et al., 2013). Plastid genome fragments were identified among the obtained contigs using the BLASTN algorithm (Altschul et al., 1990), extracted from the assembly, and utilized as “seeds” for plastid genome assembly using NOVOPlasty v3.7 (Dierckxsens et al., 2017). As an additional quality control step, completeness of the final, circularized contigs obtained from NOVOPlasty was assessed by comparison with a combination of all plastid genome contigs, extracted from each SPAdes assembly and curated using Bandage software v0.8.1 (Wick et al., 2015). Mapping of the raw reads onto the circularized contigs was performed using Bowtie2 v2.2.6 (Langmead and Salzberg, 2012) and Samtools v1.6 (Li et al., 2009); detection of putative low-coverage parts of the contigs was done using Bedtools v2.25.0 (Quinlan and Hall, 2010).

### 2.3. Plastid genome annotation and visualization

Annotation of the obtained plastid genomes was performed automatically using cpGAVAS online tool (Liu et al., 2012) and manually curated using Geneious software v10.2.6 (Kearse et al., 2012). The identity of each protein-coding gene was confirmed by cross-referencing with the NCBI non-redundant protein database using BLASTX algorithm (Altschul et al., 1990). Additionally, identification of functional domains (reverse transcriptase domain, group II intron maturase domain, and HNH endonuclease domain) within intron maturase genes, both for the four examined plastid genomes and all other euglenid plastid genomes available in the NCBI GenBank database, was performed by a HMMER algorithm-based search against the PFAM protein families’ database, using the search engine embedded within PFAM website (pfam.xfam.org; (El-Gebali et al., 2019)). Plastid genome maps were created using the OGDRAW online tool v1.3.1 (Greiner et al., 2019). Annotated plastid genomes are deposited in the public NCBI GenBank database under accession numbers OK136183 – OK136186.

### 2.4. Plastid genome-based phylogenomic analysis

Protein-coding genes of unambiguous origins (i.e., except for intron maturase genes) from the four plastid genomes of Eutreptiales investigated in this study, 29 other plastid genomes of Euglenophyta available in the NCBI GenBank database, and four plastid genomes of Pyramimonadales (Chlorophyta), were extracted from the respective plastid genomes (see Table B.1, Appendix B) and translated using Geneious software v10.2.6 (Kearse et al., 2012). Protein sequences derived from each gene were aligned using MAFFT v7.271 (Katoh and Standley, 2013), and the resulting alignments were concatenated in Geneious to produce a 37-taxa, 58-gene dataset with total length of 16,912 amino acids. The partitioned dataset was subsequently utilized as the input for maximum likelihood phylogenomic analysis using IQ-TREE software v1.6.12 (Nguyen et al., 2015), with evolution models automatically selected using *-m TEST* parameter for each partition (gene), and 1000 bootstrap replicates. Additionally, a Bayesian phylogenetic analysis on the same dataset was performed using MrBayes v3.2.6 (Ronquist et al., 2012) with 1 000 000 generations (incl. 250 000 generations burn-in), after which convergence of Markov chains was achieved.

### 2.5. Phylogenetic analysis of the plastid-encoded intron maturases

Candidate intron maturase genes (i.e., open reading frames either identified via BLAST search as homologues of previously described maturases, or found via PFAM search engine to possess any of the maturase protein domains described in section 2.3 Plastid genome annotation and visualization) were extracted from the four plastid genomes investigated in this study, as well as all 29 plastid genomes of other Euglenophyta available in the NCBI GenBank database (see Table B.1, Appendix B), and translated using Geneious software v10.2.6 (Kearse et al., 2012). The dataset was additionally complemented by a unique plastid-encoded maturase (*mat4*), identified in *Euglena geniculata* (formerly *E. myxocylindracea*), whose complete plastid genome sequence is not available (Kosmala et al., 2009; Sheveleva and Hallick, 2004).

Homologues of all 91 putative maturases were identified via BLASTP search against the NCBI non-redundant protein sequence database (NCBI-nr; (NCBI Resource Coordinators, 2018)); up to 10 best hits per maturase were extracted, deduplicated using in-house scripts, and aligned together with the euglenid plastid-encoded maturases using MAFFT v7.271 (Katoh and Standley, 2013) to produce an alignment containing 219 sequences in total. The dataset was subsequently utilized as the input for maximum likelihood phylogenetic analysis using IQ-TREE software v1.6.12 (Nguyen et al., 2015), with VT+F+I+G4 evolution model (automatically selected using *-m TEST* parameter) and 1000 bootstrap replicates.

## 3. Results and Discussion

### 3.1. Plastid genome characteristics

The general characteristics of plastid genomes of *Eutreptia* sp. CCAC1914B, *Eutreptiella* sp. CCMP389, *Eutreptiella* sp. CCMP1594 and *Eutreptiella eupharyngea* K-0027 have been shown in Table 1. Plastid genome structure of the four investigated strains have been visualized on genome maps (Figures 1a-1d). Additionally, a synteny graph of three of the analyzed genomes has been shown on Figure A.1 (Appendix A). The gene order was noticeably more similar between *Eutreptia* sp. CCAC1914B and *Etl. eupharyngea*, with only three gene block rearrangements, than between *Eutreptia* sp. and *Eutreptiella* sp. CCMP1594 (seven rearrangements) or *Etl. eupharyngea* and *Eutreptiella* sp. CCMP1594 (nine rearrangements).

**Table 1.**
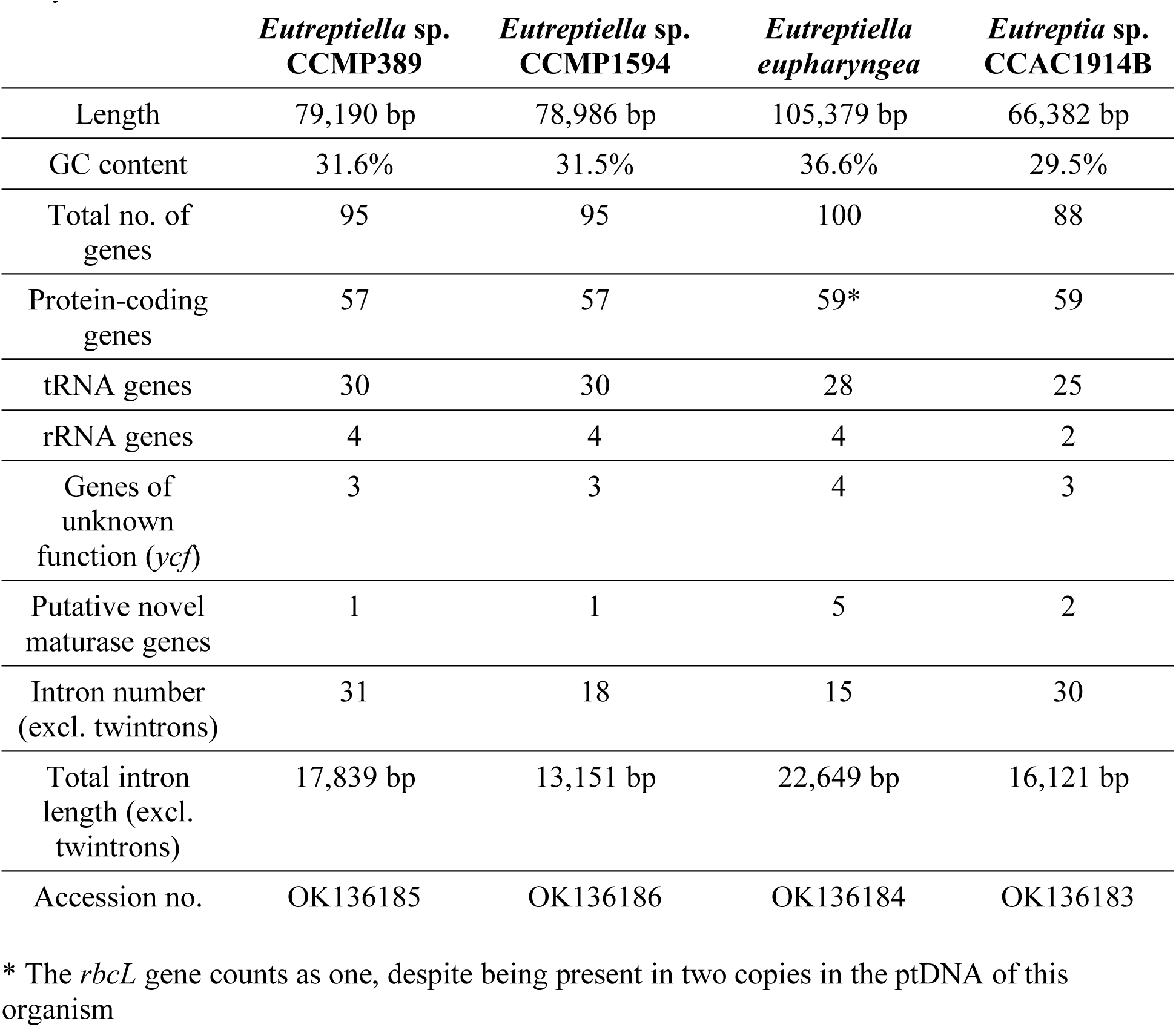
Characteristics of the four new plastid genomes of Eutreptiales presented in this study.

|  | *Eutreptiella* sp.<br>CCMP389 | *Eutreptiella* sp.<br>CCMP1594 | *Eutreptiella*<br>*eupharyngea* | *Eutreptia* sp.<br>CCAC1914B |
| --- | --- | --- | --- | --- |
| Length | 79,190 bp | 78,986 bp | 105,379 bp | 66,382 bp |
| GC content | 31.6% | 31.5% | 36.6% | 29.5% |
| Total no. of genes | 95 | 95 | 100 | 88 |
| Protein-coding genes | 57 | 57 | 59* | 59 |
| tRNA genes | 30 | 30 | 28 | 25 |
| rRNA genes | 4 | 4 | 4 | 2 |
| Genes of unknown function (*ycf*) | 3 | 3 | 4 | 3 |
| Putative novel maturase genes | 1 | 1 | 5 | 2 |
| Intron number (excl. twintrons) | 31 | 18 | 15 | 30 |
| Total intron length (excl. twintrons) | 17,839 bp | 13,151 bp | 22,649 bp | 16,121 bp |
| Accession no. | OK136185 | OK136186 | OK136184 | OK136183 |

\* The *rbcL* gene counts as one, despite being present in two copies in the ptDNA of this organism

**Figure 1.**
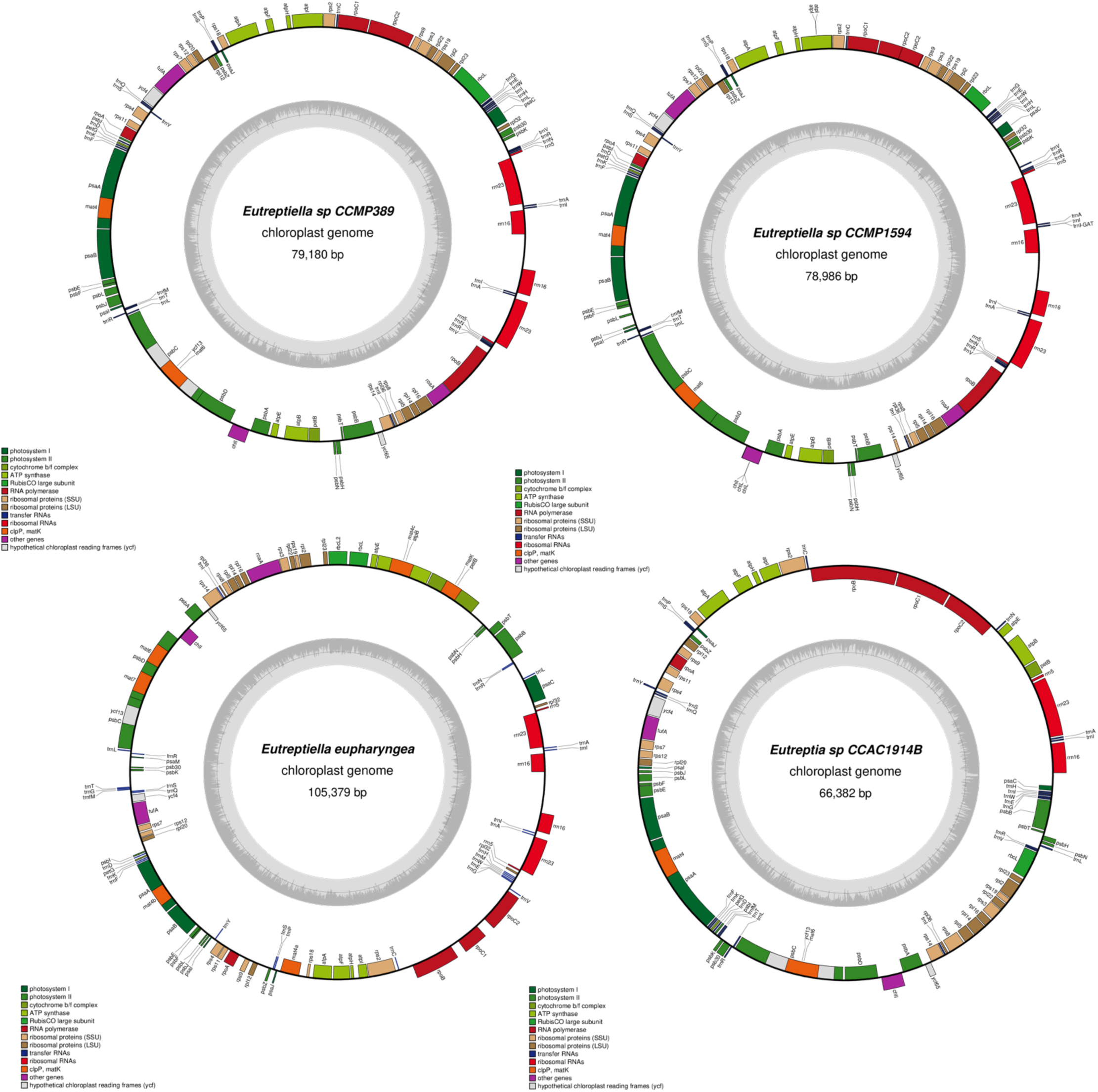
Chloroplast genome maps of *Eutreptiella* sp. CCMP389 (A), *Eutreptiella* sp. CCMP1594 (B), *Eutreptiella eupharyngea* K-0027 (C) and *Eutreptia* sp. CCAC1914B (D). Maps were generated using OGDraw online tool v1.3.1 (Greiner et al., 2019).

The four plastid genomes are noticeably different in size, which ranges from approximately 66 kbp (*Eutreptia* sp.) to over 105 kbp (*Etl. eupharyngea*). This variability, however, does not reflect the differences in coding contents of these genomes, but instead results from differences in the quantities of non-coding and repeated sequences, which has been previously observed in comparative analyses of plastid genomes of freshwater euglenids (Karnkowska et al., 2018). The protein-coding gene complements of the analyzed genomes, while being largely similar, seem to bear only one substantial difference (except for the intron-encoded protein content, which will be described in further detail in separate sections): the gene *psaM*, encoding a small photosystem I subunit, was successfully identified only in *Etl. eupharyngea*. As this particular gene was also reportedly missing in the plastid genome of *Eutreptia viridis* (Wiegert et al., 2012), its absence in some other Eutreptiales is not surprising.

Although *Eutreptiella* sp. CCMP389 and *Eutreptiella* sp. CCMP1594 seem to have nearly identical plastid genomes, as indicated by their very similar size and identical gene complement, their intron repertoire is noticeably different, both in terms of intron number and their total length (Table 1). Moreover, although introns are more numerous in both of the two than in *Etl. eupharyngea*, it is the latter that has greater total intron length.

Despite the photosynthetic euglenids being known to exhibit a vast variety of rDNA operon organization types (Karnkowska et al., 2018), all three new plastid genomes of *Eutreptiella* spp. possess a pair of inverted rDNA repeats, separated by a short (ca. 2-3 kbp-long), entirely non-coding small single-copy region. In contrast, the investigated *Eutreptia* sp. strain turned out to possess a single copy of the ribosomal operon, similarly to *Eutreptia viridis –* the only other representative of the genus with available complete plastid genome sequence (Wiegert et al., 2012).

### 3.2. Polyphyly of the genus *Eutreptiella*

The plastid gene complement-based phylogenomic tree of euglenophytes has been shown on Figure 2. Although Euglenales have been recovered as a well-supported monophyletic group, similarly to previous plastid-based phylogeny (Karnkowska et al., 2018), we observed that Eutreptiales are paraphyletic with respect to Euglenales and divided into two clades: *Eutreptia* spp. + *Eutreptiella* sp. CCMP389 + *Eutreptiella* sp. CCMP1594 form a sister clade to freshwater euglenids, while *Etl. eupharyngea + Etl. pomquetensis* + *Etl. gymnastica* branch off earlier (all relationships recovered with absolute bootstrap support). For the sake of clarity, the terms *Eutreptiella* clade I (synonymous with “*Eutreptiella* clade” in (Kolisko et al., 2020), encompassing *Eutreptiella* sp. CCMP389 and *Eutreptiella* sp. CCMP1594) and clade II (a collective name for both “*Eutreptiella eupharyngea* clade” and “*Eutreptiella gymnastica* clade” in (Kolisko et al., 2020), encompassing *Etl. eupharyngea, Etl. pomquetensis* and *Etl. gymnastica*) will be used in this paper to distinguish between the *Eutreptiella* strains that form the proximal branch to *Eutreptia*, and those which form a branch at the base of all photosynthetic euglenids, respectively.

**Figure 2.**
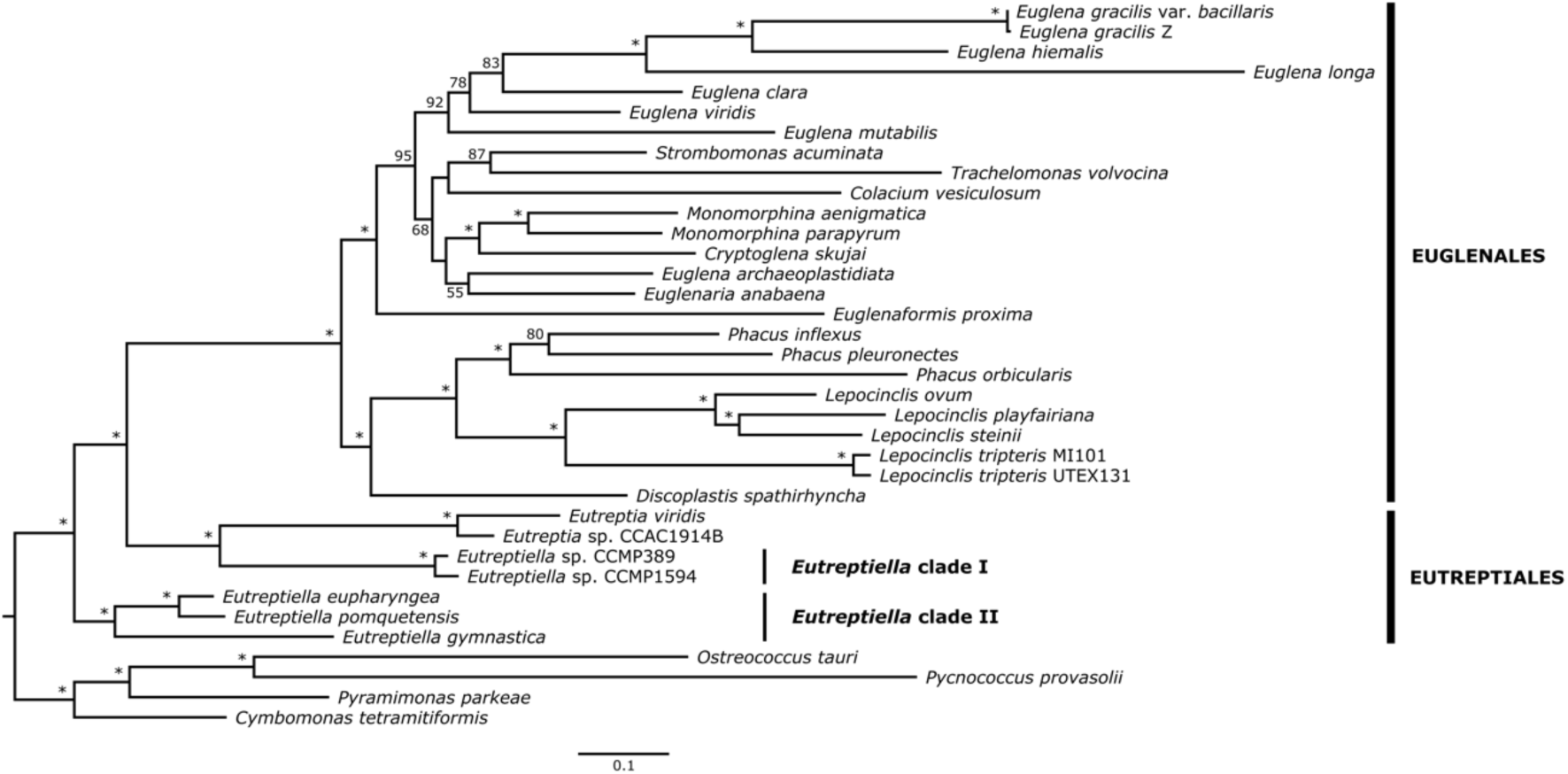
Plastid-based maximum likelihood phylogenetic tree of Euglenophyta. Nodes with absolute support values are marked with asterisks (*). Support values below 50 are not shown.

A vast array of previous works has shown that there is a general incongruence in the postulated phylogenetic relationships within the Eutreptiales group (Bicudo and Menezes, 2016; Karnkowska et al., 2018; Kolisko et al., 2020; Marin et al., 2003; Yamaguchi et al., 2012). Moreover, these disparities cannot be attributed to differences in selected molecular markers (encompassing nuclear and plastid-encoded rDNA and protein-coding genes), as even phylogenies based on the same marker – most commonly nuclear SSU rDNA – can have different topologies (Kolisko et al., 2020; Marin et al., 2003; Yamaguchi et al., 2012). By using diverse datasets, including both different markers and different representatives of marine euglenids’ taxa, the genus *Eutreptiella* has been recovered either as monophyletic (Karnkowska et al., 2018; Kolisko et al., 2020; Yamaguchi et al., 2012) or as paraphyletic with respect to *Eutreptia* (Bicudo and Menezes, 2016; Karnkowska et al., 2015; Marin et al., 2003), but the support for both possibilities was highly variable. Similar disparities were observed in the recovery of the Eutreptiales group, either as monophyletic (Bicudo and Menezes, 2016; Karnkowska et al., 2015; Kolisko et al., 2020; Yamaguchi et al., 2012), or as paraphyletic with respect to Euglenales (Karnkowska et al., 2018; Marin et al., 2003).

Unfortunately, the two most recent euglenophyte phylogenies were vastly different in terms of input dataset construction methodology – the plastid-based phylogenomic analysis by Karnkowska et al. 2018 had a very scarce representation of marine euglenids, as it the study focused on freshwater taxa, while the phylogeny of Kolisko et al. 2020 used a broadly-sampled single marker gene – nuclear SSU rDNA. As a result, it is difficult to compare the validity of findings based on those two datasets. Considering that this work has doubled the available plastid-derived genomic data complement of Eutreptiales, enabling us to construct a comprehensive dataset, we decided to utilize it to obtain the best currently attainable resolution of this group’s phylogeny.

Quite surprisingly, the results of our maximum-likelihood phylogenomic analysis are not fully congruent with any phylogeny reconstructed before, with both Eutreptiales and the genus *Eutreptiella* being polyphyletic (Figure 2). They are, however, congruent in all nodes relevant to this work with the Bayesian phylogeny reconstruction we performed using the same input dataset (Figure A.2, Appendix A). As a side note, incongruencies between our ML tree and Bayesian tree occurred only within the Euglenales clade, with divergent positions and contents of its subordinate clades, which are not the subject of this study. Moreover, the phylogenetic relationships between the two families within Euglenales and almost all of their subordinate genera (with a few exceptional *incertae sedis* taxa, e.g., *Euglena archaeoplastidiata* or *Euglenaria* spp.) have been resolved by numerous congruent, robustly supported multigene phylogenies, presented in other studies (Karnkowska et al., 2018, 2015; Kim et al., 2015).

Interestingly, although both Eutreptiales and *Eutreptiella* were resolved as polyphyletic taxa only by Marin et al., the phylogeny proposed in that study had “inverted” topology within Eutreptiales – the *Eutreptia* + *Eutreptiella* sp. CCMP389 clade was at the base of all photosynthetic euglenids, while the clade encompassing all other *Eutreptiella* strains was the sister one to Euglenales (Marin et al., 2003).

### 3.3. Novel maturases in *Eutreptiella* clade II and other euglenids

The key difference between protein-coding gene complements of *Eutreptiella* clades I & II lies in the number and location of group II intron maturases. Apart from the previously described *mat1* (*ycf13*), present in a *psbC* gene intron in all plastid genomes of euglenids except *Euglena longa*, and the patchily-distributed *mat2* (encoded within a different *psbC* intron) and *mat5* (encoded within *psbA* intron), we discovered that the ptDNA of *Etl. eupharyngea* carries six additional maturase genes. Among those, we identified one gene to be a homologue of *matK* – a maturase commonly found in plant plastids (Barthet et al., 2020). However, the origins of the remaining five were not as clear, and unraveling them required a phylogenetic analysis.

As shown on the phylogenetic tree of euglenid plastid-encoded maturases (Figure 3; Figure A.3, Appendix A), three of the putative maturase genes from *Etl. eupharyngea* turned out to share their origins with *mat4* – a gene previously identified, among euglenids, only in *Euglena geniculata* (Sheveleva and Hallick, 2004). These three putative maturases will be further referred to as *mat4a* (encoded as a regular ORF, not within an intron), *mat4b* (encoded as an IEP within *psaA* gene intron) and *mat4c* (encoded within *atpB* intron). The other two of five, however, do not seem to be homologous to any specific known maturase genes, and therefore, we decided to assign new names to them: *mat6* (encoded as an IEP within *psbD* intron 1) and *mat7* (encoded within *psbD* intron 2). Additionally, we identified four hypothetical proteins, described previously in *Etl. pomquetensis* (Dabbagh et al., 2017), to be close homologues of *mat4abc* and *mat6*.

**Figure 3.**
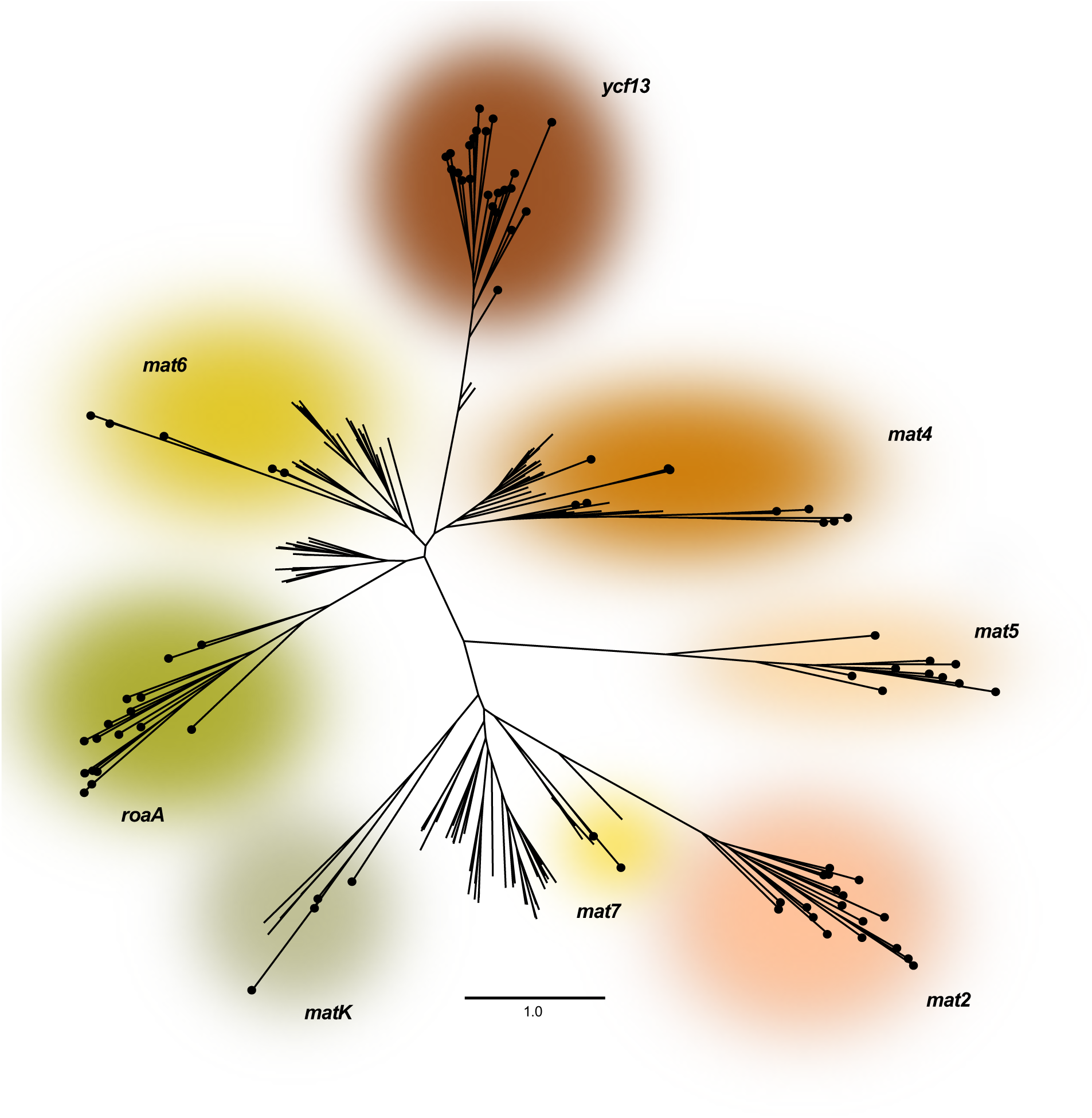
Schematic phylogenetic tree of euglenid plastid-encoded intron maturases and their known homologs. Distinct colors denote orthologue groups of euglenid plastid-derived sequences. Black dots on branch tips denote sequences extracted from euglenid plastid genomes.

Based on these findings, we decided to broaden the scope of our maturase survey to include all available plastid genomes of Euglenophyta, and found seven genes of previously unknown function to exhibit detectable homology with maturase genes. These include: a putative maturase gene encoded within the first *rbcL* gene intron in *Discoplastis spathirhyncha* (homologous to *mat6*), one putative maturase encoded within an *rbcL* intron in *Cryptoglena skujai* (homologous to *mat6*), *Euglena archaeoplastidiata* (homologous to *mat7*) and *Euglenaria anabaena* (homologous to *mat7*), one putative maturase encoded within a *psbC* intron in *Lepocinclis steinii* (homologous to *mat7*), one putative maturase encoded within a *psbA* intron in *Trachelomonas volvocina* (homologous to *matK*), and one putative non-intron-encoded maturase in *Eutreptiella gymnastica* (homologous to *mat5*). The full results of our search are depicted on Figure 4. Given that group II intron maturases have been identified in introns within tRNA genes in plastid genomes of rhodophytes (Janouškovec et al., 2013) and land plants (Hausner et al., 2006), we additionally screened the available euglenid genomic data for IEPs within tRNA gene introns. Despite the abundance of introns and the great variety of locations of intron maturases in euglenid plastids, we found no tRNA gene introns in euglenid plastid genomes at all.

**Figure 4.**
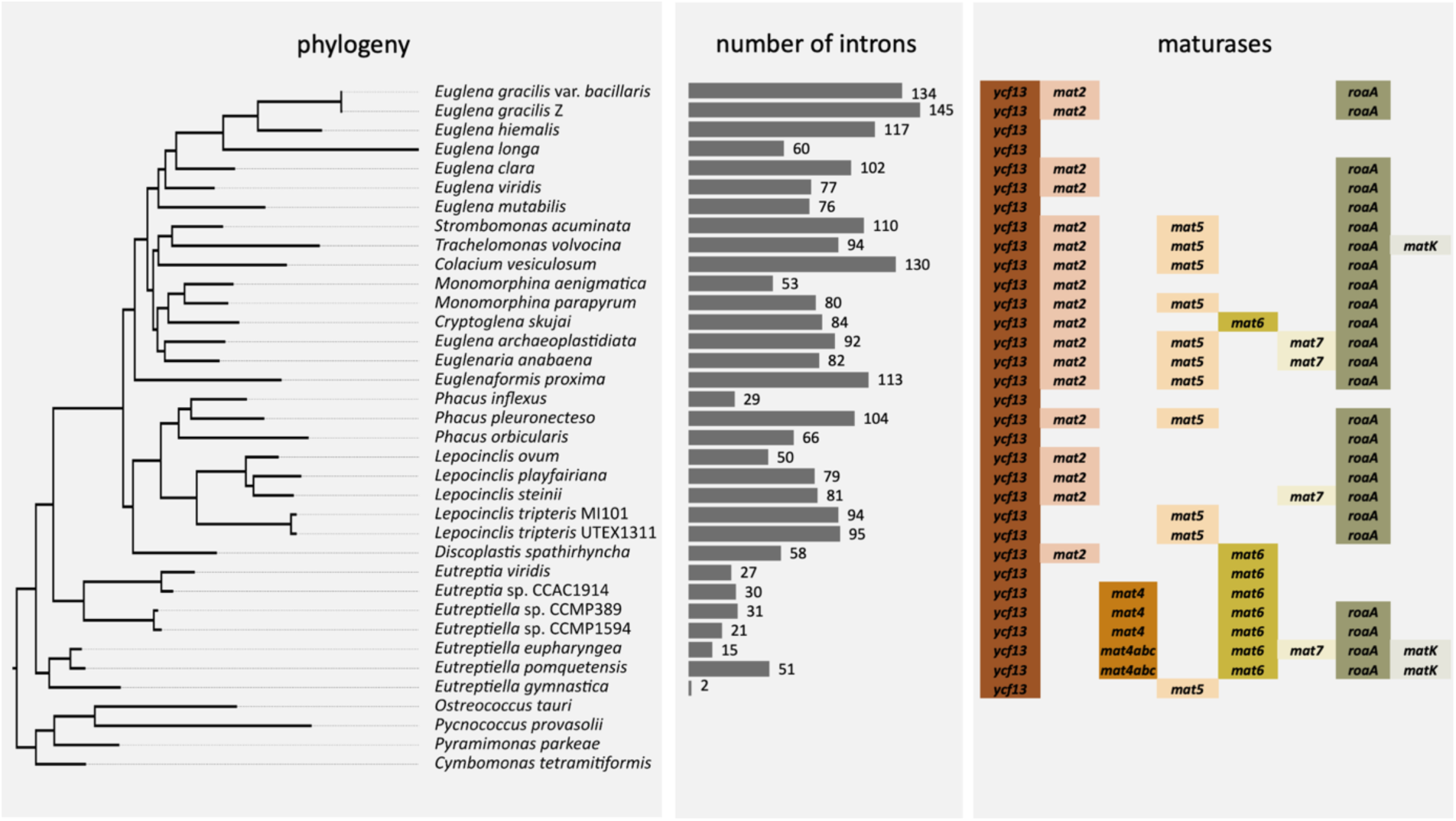
Numbers of introns and presence of maturases identified in plastid genomes of Euglenophyta, mapped onto the group’s phylogeny.

### 3.4. *Eutreptia* and *Eutreptiella* clade I plastid genomes share a unique trait: a maturase encoded within another maturase

In a previous study, it has been shown that the *ycf13* gene of *Eutreptia viridis*, while being an IEP itself, possesses a hypothetical protein-coding ORF within its second intron (Wiegert et al., 2012). This finding, however interesting, has been commented upon rather sparsely in the study itself, and the possible character of this hypothetical protein has not been indicated. Considering our aforementioned findings that multiple hypothetical proteins in euglenid plastids may, in fact, be novel intron maturases, we decided to thoroughly investigate the *ycf13* gene’s organization in the new plastid genomes of Eutreptiales. As a result, we identified the presence of an undoubtedly orthologous ORF within the second intron of the *ycf13* gene in the plastid genomes of *Eutreptia* sp. CCAC1914B, *Eutreptiella* sp. CCMP389 and *Eutreptiella* sp. CCMP1594, but not in any other euglenid, including *Etl. eupharyngea*. Moreover, a HMMER search revealed all four ORFs (including the one from *Eut. viridis*) to contain a group II intron maturase domain, thus suggesting that they constitute an orthologous group of yet another previously undescribed plastid-encoded maturase. The following phylogenetic analysis revealed the *ycf13* intron-encoded maturases to be, in fact, homologues of *mat6*.

This curiosity is, to our knowledge, the first documented case of a “three-storeyed protein”, i.e., a gene located in an intron, in a gene within an intron, within yet another gene. In the specific context of the evolution of plastid genomes in Euglenophyta, it is also a strong indicator towards the presence of a timeline, and not a single burst, of acquisition of group II introns and their maturases; even without any knowledge of the precise time scale, the embedding of an IEP within another IEP naturally implies the second one to be older. In addition, it is also the first known case of a pair of genes forming a true “matryoshka”, since *mat6* and *ycf13* have common evolutionary origins, the same (at least ancestrally) biological functions, and are located one within another.

It is, however, rather puzzling whether this exceptionally convoluted genetic setting serves any specific biological function. The most similar structure described so far (although one “storey” shorter) had been discovered in the mitochondrial genome of an ascomycete fungus *Chaetomium thermophilum*, and consists of an internal group II intron within a homing endonuclease gene, encoded within an external group I intron in the small ribosomal subunit gene (Guha and Hausner, 2014). The authors of the study suggested that the excision of the internal intron might be an “on” switch for the homing endonuclease (Guha and Hausner, 2014); however, in case of *mat6/ycf13*, the presence of the innermost gene (*mat6*) would be likely redundant as a part of the switching mechanism of *ycf13*. Unfortunately, without detailed *in vivo* studies, it can only be speculated if *mat6* and *ycf13* are in any way functionally connected (e.g., whether the product of *mat6* aids the excision of its outer introns, or if a viable hybrid intron maturase is produced if the innermost intron is not excised), or if their topological relationship is merely the result of a very odd coincidence.

### 3.5. *roaA* is an intron maturase-derived pseudogene

Our search for new maturase genes has also yielded another interesting result: we identified a long-known, common plastid-encoded gene to be, in fact, yet another maturase. We found an additional putative maturase, initially labeled *mat8*, in the ptDNA of *Etl. eupharyngea*, but a more thorough search revealed it to be, most likely, a homologue of *roaA*. Despite being present in a vast majority of euglenid plastid genomes, very little is known about the activity and function of this particular gene. In fact, the only occurrence of this gene’s name in literature is a study of *E. gracilis*’s *roaA*, which revealed this gene to be co-transcribed with a number of other ribosomal proteins (hence its name – an abbreviation of “ribosomal operon-associated protein A”) and a possible RNA-binding capability, deduced from the analysis of its amino acid sequence (Jenkins et al., 1995). These observations led us to suspect that *roaA* might be a maturase itself, and a HMMER-based protein domain survey of all euglenid *roaA* sequences not only confirmed our suspicions, but also revealed an complicated pattern of reductive evolution of this gene in Euglenophyta (Figure 4).

We observed that euglenid *roaA* genes can carry the reverse transcriptase domain (RVT) and group II intron maturase domain (GIIM), but both of them have been differentially lost a number of times – in fact, the only organism to possess the “ancestrally functional” (i.e., possessing both the RVT and the GIIM domain) *roaA* is *Etl. pomquetensis* (Table 2). At the same time, *roaA* genes in most euglenids (e.g. *Lepocinclis* spp., *Monomorphina spp*., *Phacus pleuronectes* or *Euglena viridis*) have only retained the GIIM domain, while in *Etl. eupharyngea* and *Eutreptiella* sp. CCMP1594, the RVT domain has been retained instead. What is more, in some other euglenophytes, such as *Euglena clara, Colacium vesiculosum* or *Eutreptiella* sp. CCMP389, *roaA* has not retained any of the two domains, thus having no discernible biological functionality.

**Table 2.** Conserved protein domains in euglenid plastid-encoded maturases. RVT = reverse transcriptase (RNA-dependent DNA polymerase) domain. GIIM = group II intron maturase domain. HNHE = HNH endonuclease domain.

| **Organism, gene name and/or location** | **Conserved domains** |
| --- | --- |
| *C. skujai mat2* | - |
| *C. skujai mat6 (rbcL intron)* | RVT + GIIM + HNHE |
| *C. skujai roaA* | GIIM |
| *C. skujai ycf13* | RVT |
| *Co. vesiculosum mat2* | GIIM |
| *Co. vesiculosum mat5* | - |
| *Co. vesiculosum roaA* | - |
| *Co. vesiculosum ycf13* | - |
| *D. spathirhyncha mat2* | - |
| *D. spathirhyncha mat6 (rbcL intron)* | RVT + GIIM + HNHE |
| *D. spathirhyncha ycf13* | RVT |
| *E. archaeoplastidiata mat2* | - |
| *E. archaeoplastidiata mat5* | - |
| *E. archaeoplastidiata mat7 (rbcL intron)* | GIIM |
| *E. archaeoplastidiata roaA* | GIIM |
| *E. archaeoplastidiata ycf13* | RVT |
| *E. clara mat2* | - |
| *E. clara roaA* | - |
| *E. clara ycf13* | RVT |
| *E. gracilis* var. *bacillaris* *mat2* | - |
| *E. gracilis* var. *bacillaris* *roaA* | - |
| *E. gracilis* var. *bacillaris* *ycf13* | RVT |
| *E. gracilis* Z *ycf13* | RVT |
| *E. hiemalis* *ycf13* | RVT |
| *E. longa* *ycf13* | RVT |
| *E. mutabilis* *roaA* | - |
| *E. mutabilis* *ycf13* | RVT |
| *E. myxocylindracea* *mat4* | RVT + GIIM + HNHE |
| *E. viridis* *mat2* | - |
| *E. viridis* *roaA* | GIIM |
| *E. viridis* *ycf13* | RVT |
| *Ef. proxima* *mat2* | GIIM |
| *Ef. proxima* *mat5* | - |
| *Ef. proxima* *roaA* | GIIM |
| *Ef. proxima* *ycf13* | RVT |
| *Er. anabaena* *mat2* | - |
| *Er. anabaena* *mat5* | - |
| *Er. anabaena mat7 (rbcL intron)* | RVT + GIIM |
| *Er. anabaena roaA* | GIIM |
| *Er. anabaena ycf13* | RVT |
| *Etl. eupharyngea mat4a* | RVT + GIIM |
| *Etl. eupharyngea mat4b* | RVT + GIIM |
| *Etl. eupharyngea mat4c* | RVT + GIIM + HNHE |
| *Etl. eupharyngea mat6* | GIIM |
| *Etl. eupharyngea mat7* | RVT + GIIM |
| *Etl. eupharyngea matK* | GIIM |
| *Etl. eupharyngea roaA* | RVT |
| *Etl. eupharyngea ycf13* | RVT |
| *Etl. gymnastica mat5* | - |
| *Etl. pomquetensis mat4a* | RVT + GIIM |
| *Etl. pomquetensis mat4b* | RVT + GIIM |
| *Etl. pomquetensis mat4c* | RVT + GIIM + HNHE |
| *Etl. pomquetensis mat6* | GIIM |
| *Etl. pomquetensis matK* | GIIM |
| *Etl. pomquetensis roaA* | RVT + GIIM |
| *Etl. pomquetensis ycf13* | RVT |
| *Eut. viridis mat6 (ycf13 intron)* | GIIM |
| *Eut. viridis ycf13* | RVT + GIIM |
| *Eutreptiella* sp. CCMP1594 *mat6* (*ycf13* intron) | RVT + GIIM |
| *Eutreptiella* sp. CCMP1594 *roaA* | RVT |
| *Eutreptiella* sp. CCMP1594 *ycf13* | RVT |
| *Eutreptiella* sp. CCMP389 *roaA* | - |
| *Eutreptiella* sp. CCMP389 *ycf13* | RVT + GIIM |
| *L. ovum mat2* | - |
| *L. ovum roaA* | GIIM |
| *L. ovum ycf13* | RVT |
| *L. playfairiana mat2* | GIIM |
| *L. playfairiana roaA* | GIIM |
| *L. playfairiana ycf13* | RVT |
| *L. steinii mat2* | - |
| *L. steinii mat7* (*psbC* intron) | RVT + GIIM |
| *L. steinii roaA* | GIIM |
| *L. steinii ycf13* | RVT |
| *L. tripteris* MI101 *mat5* | - |
| *L. tripteris* MI101 *roaA* | GIIM |
| *L. tripteris* MI101 *ycf13* | RVT |
| *L. tripteris* UTEX1311 *mat5* | - |
| *L. tripteris* UTEX1311 *roaA* | GIIM |
| *L. tripteris* UTEX1311 *ycf13* | RVT |
| *M. aenigmatica* *mat2* | - |
| *M. aenigmatica* *roaA* | GIIM |
| *M. parapyrum* *mat2* | - |
| *M. parapyrum* *mat5* | - |
| *M. parapyrum* *roaA* | GIIM |
| *M. parapyrum* *ycf13* | RVT |
| *Ph. inflexus* *ycf13* | RVT |
| *Ph. orbicularis* *roaA* | - |
| *Ph. orbicularis* *ycf13* | RVT |
| *Ph. pleuronectes* *mat2* | - |
| *Ph. pleuronectes* *mat5* | - |
| *Ph. pleuronectes* *roaA* | GIIM |
| *Ph. pleuronectes* *ycf13* | RVT |
| *S. acuminata* *mat2* | - |
| *S. acuminata* *mat5* | - |
| *S. acuminata* *roaA* | GIIM |
| *S. acuminata* *ycf13* | RVT |
| *T. volvocina* *mat5* | - |
| *T. volvocina matK* (*psbA* intron) | RVT + GIIM |
| *T. volvocina roaA* | GIIM |
| *T. volvocina ycf13* | RVT |
| *T. volvocina mat2* | GIIM |

As evidenced above, the domain loss pattern in euglenid *roaA* seems random, as it is not in any way consistent with the phylogenetic relationships between organisms, therefore indicating very low, if any at all, selective pressure for retention of any of its functional domains. Additionally, euglenid *roaA* amino acid sequences are highly divergent – the protein sequence similarity between *roaA* of different species, even within one genus, is exceptionally low (ca. 30%). This leads us to speculate that this gene might, as a matter of fact, have undergone transition to being a pseudogene early in the evolution of euglenophytes (or even before the plastid acquisition itself), and it serves no biological function whatsoever in the contemporary euglenid plastids.

### 3.6. Conventional euglenid plastid-encoded maturases are functionally deficient

Considering our aforementioned observations on functional reduction in certain maturase genes in euglenid ptDNA, we screened all other euglenid maturases, both described in past studies and newly-found, for the presence of maturase protein domains in order to verify their functional capabilities. The results of this analysis are shown in Table 2.

Based on the results of our analysis, it becomes quite clear that functional reduction is not limited to the *roaA* family, but it is in fact prevalent among plastid-encoded maturases in euglenids. The maturase *ycf13 (mat1)*, identified in almost all euglenid plastid genomes, turned out to possess only the RVT domain, with the sole exception of *Eutreptia viridis ycf13*, which carries the GIIM domain as well, indicating that it is not, from a functional point of view, a maturase, and it has no capability of aiding the excision of introns in euglenid ptDNA. What function could a reverse transcriptase have in a plastid genome, however, remains a mystery, but the retention of this activity in all *ycf13* proteins in euglenids (except the one in *Colacium vesiculosum*) indicates that it still serves a purpose. Interestingly, although gradual domain loss is rather frequent in known IEPs and is considered to be indicative of their degenerative evolutionary path, the GIIM domain is always the last one to be retained, not RVT (Zimmerly and Semper, 2015).

Astonishingly, the reduction of other conventional euglenid maturases, *mat2* and *mat5*, is even more drastic. Out of 17 *mat2* orthologs analyzed, we found only four (extracted from the ptDNA of *Colacium vesiculosum, Euglenaformis proxima, Trachelomonas volvocina* and *Lepocinclis playfairiana*) to retain a functional GIIM domain, while the remaining 13 possess no functional protein domain at all, indicating complete loss of function. What is more, none of the 10 *mat5* orthologs contains any functional domain as well. Therefore, unless plastid-encoded maturases of euglenids are so divergent that the domains responsible for their intron-related activity cannot be detected by homology-based search methods, it would seem evident that plastid genomes of most euglenophytes, despite being highly intron-rich, do not encode any functional intron maturases. These findings are congruent with the previous observations of widespread evolutionary “decay” of intron structure and catalytic activity of their RNA components in euglenid plastids (Lambowitz and Belfort, 2015); nonetheless, it is rather astounding that these maturases, despite being prevalently deprived of function, have almost invariantly retained the capability for expression as intact ORFs, with no start codon losses, stop codon insertions or frameshifts.

The functional deficiency of the investigated plastid-encoded intron maturases further supports the idea that intron excision from plastid-encoded genes in euglenids must be supported by a different mechanism providing maturase-like activity *in trans*, possibly a nuclear-encoded one, as it has been documented in organellar (primarily mitochondrial) introns of other eukaryotic groups (Zimmerly and Semper, 2015). Unfortunately, with nuclear genome data from photosynthetic euglenids being extremely scarce, this hypothesis would be difficult to verify at this point; we did, however, identify the sequence *EG_transcript_10014* in the published transcriptome of *Euglena gracilis* as a functional nuclear maturase (Ebenezer et al., 2019).

It is also interesting that among all euglenid plastid-encoded maturases subjected to the aforementioned analysis, the only ones to possess all conventional maturase domains, i.e. RVT, GIIM (“X”) and HNH endonuclease domain (HNHE/”En”), are the *mat4* of *E. geniculata* and four novel maturases identified in this study – the *mat4c* maturases of *Etl. eupharyngea* and *Etl. pomquetensis*, and the *mat6* maturases of *Cryptoglena skujai* and *Discoplastis spathirhyncha* (Table 2; Figure A.4, Appendix A). These five maturases most likely possess the capabilities not only for group II intron excision, but also retromobility – a trait which has been frequently lost in non-bacterial intron maturases (Zimmerly and Semper, 2015).

On the other hand, the overwhelming prevalence of functionally-deficient maturases over intact ones in euglenid plastids should not come as a surprise, considering that all group II introns used to contain a functional maturase ORF at some point in their evolutionary past (Dai and Zimmerly, 2002). It would therefore seem that the proposed intron “life cycle” (which consists of: gain via retrotransposition, ORF degeneration with retained splicing capability, ORF loss, and secondary deletion of the entire intron (Dai and Zimmerly, 2002)) is impaired in euglenid plastids due to a bottleneck in the last stage, responsible for excessive accumulation of intron “husks”, deprived of their ORFs. Bearing in mind that this situation has arisen in a secondary endosymbiosis-derived plastid, this bottleneck might have been caused by incomplete inheritance of the organellar intron maintenance mechanism by its new host.

It is also worth noting that with all new available data, there is hardly any convincing argument for the previously proposed hypothesis of a positive correlation between the number of introns and the number of intron maturase genes within a plastid genome (Karnkowska et al., 2018). Not only is there little biological basis for such correlation to occur, with the vast majority of maturases lacking key functional domains, but it is also not supported statistically (Pearson’s product moment correlation; *df* = 29, *r* = 0.097). Still, we believe that the key to unraveling the true capabilities of euglenid plastid-encoded maturases might lie in further *in vivo* studies, especially considering that such studies on this subject in the past, while rare, have been rather illuminating, providing key insights into the specificity of maturases’ activity, which could not be studied via *in silico* analyses (Barthet et al., 2020; Sheveleva and Hallick, 2004).

### 3.7. Mosaic-like origins of euglenid plastid-encoded maturases

Our phylogenetic analysis of the euglenid plastid-encoded putative maturases (Figure 3) revealed that while all ortholog groups proposed to date form rather well-supported clades, the closest relatives for each of these clades are highly diverse. *ycf13* and *mat4* branch within groups of cyanobacterial and chlorophytean reverse transcriptases/maturases (partially in congruence with the ancestry of *mat4* from *E. geniculata*, proposed by Sheveleva and Hallick (Sheveleva and Hallick, 2004)), suggesting their transfer into the ancestor of euglenophytes along with the secondary plastid. Nonetheless, the clades encompassing them are clearly separate, indicating that these maturases may have originated by paralogous duplication in chlorophytes before the secondary plastid endosymbiosis, or possibly even in cyanobacteria, before the first plastid came into existence.

The hypothesis of the early divergence of these maturases is further supported by their phylogenetic vicinity to different green algae – *ycf13* branch contains a putative plastid maturase ORF from a pyramimonadalean, *Pyramimonas parkeae* (a postulated closest extant relative of the euglenid plastid), while *mat4* is clustered together with genes from mamiellophytes (*Ostreococcus tauri*) and ulvophytes (*Derbesia* sp., *Pseudoneochloris marina*) (Figure 3). This also indicates that the green alga which became the photosynthetic symbiont of euglenids most likely carried both *ycf13* and *mat4*. Such combination has not been found in any extant chlorophytes, suggesting that the kinship between known green algal species and the euglenid plastids’ ancestor may not be as close as previously thought.

The origins of *mat7* and *matK* in euglenophytes are likely similar, as both of their ortholog groups form clades with green algal maturases. Interestingly, the evolutionary history of *mat2* is slightly more complex – its orthogroup’s close phylogenetic relationship with plant mitochondrial maturases suggests that this gene had been originally mitochondrial in Archaeplastida, and it had probably undergone transfer to the plastid genome before being acquired by euglenophytes via secondary plastid endosymbiosis.

On the other hand, the phylogenetic positions of other euglenid maturases are indicative of horizontal gene transfers from a variety of donors. The *mat6* gene cluster branches within a clade of reverse transcriptases/maturases of Firmicutes, while the clade encompassing *roaA* orthologs contains genes from diverse prokaryotes, such as *Streptococcus* (Firmicutes), *Escherichia coli* (Gammaproteobacteria), and even *Methanosarcina* (Euryarchaeota). Additionally, the *mat5* cluster is located in a rather puzzling position at the base of a large portion of the tree, encompassing most of the aforementioned clades, with no immediate neighbors. With very low sequence similarity and support values for these clades, it is difficult to conclude when, how, and from whom were these maturases transferred to the euglenids’ plastid genomes. However, given how widespread all of them are across the phylogenetic tree of Euglenophyta – except for the freshwater-exclusive *mat2*, every maturase gene has been found both in Euglenales and Eutreptiales – it is almost certain that their acquisition occurred in the last common ancestor of plastid-bearing euglenids or earlier.

It is also worth noting that this phylogenetic analysis of euglenid maturases might have revealed an additional case of acquisition of a maturase gene outside of euglenophytes, in a secondary red plastid lineage. We identified that a unique IEP of *Dictyocha speculum* (Dictyochophyceae, SAR), encoded within the only intron ever found among all six available plastid genomes of this group (Han et al., 2019; Kayama et al., 2020), is in fact a *mat4* homologue, containing all three functional protein domains found in *mat4c* of *Etl. eupharyngea* and *Etl. pomquetensis*. The sisterhood of *D. speculum*’s *mat4* and a putative maturase of *Corynoplastis japonica* (Rhodophyta) may indicate that the maturase has been transferred to the dictyochophyte along with its secondary red alga-derived plastid; however, as this is the only homolog of this particular maturase among all ochrophytean plastid genomes, the possibility of a horizontal transfer of this gene independently from the plastid endosymbiosis cannot be excluded.

Additionally, we were quite surprised to find that, in contrary to previous findings (Karnkowska et al., 2018), euglenid plastid-encoded maturases do not have strictly conserved locations within specific introns of other genes. Although the “well-known” maturases, such as *ycf13* or *mat2*, are indeed located in corresponding intron sites across all euglenophytes (e.g. *ycf13* within the *psbC* intron), this is not the case with the newly-identified maturases, such as *mat6*, located in *psbD* intron in *Etl. eupharyngea*, in *rbcL* intron in *Discoplastis spathirhyncha*, and in *psbC/ycf13* twintron in *Eutreptiella* sp. CCMP389. Therefore, maturases within one orthologous group might either be the products of a single transfer to the plastid genome with subsequent duplication, probably via intragenomic homing (Sheveleva and Hallick, 2004), and differential loss, or the results of multiple independent transfers, possibly from the plastid host’s nucleus. It has been documented that a nuclear-encoded maturase can successfully operate *in trans* in organellar-encoded gene introns, e.g. in plant mitochondria (Brown et al., 2014), however, repeated transfers of maturase genes to new sites within the organellar genome may be inevitable, considering that facilitation of own horizontal transfer is among their core functions.

## 4. Conclusions

In this study, we obtained four new plastid genomes of marine euglenids. Combined with almost 30 other plastid genomes of euglenophytes available to date, we provided a comprehensive dataset to illuminate the phylogeny of this group and the evolution of its organellar genomes. The phylogenetic relationships between the representatives of the marine clades turned out quite differently than expected and indicated both the genus Eutreptiella and order Eutreptiales to be paraphyletic. However, the most interesting and surprising results came from the analysis of group II intron maturase genes. We did not only find new orthologues of previously described genes in new species or new locations within ptDNA, but we also uncovered two completely new genes with distinct phylogenetic distribution and evolutionary history. In all cases, however, the retained biological functionality within a maturase orthologue cluster varies with the species or strain, implying a frequent inactivation and eventual loss of maturases over time. The phylogenetic analyses of maturases also highlighted their diverse origins, encompassing genes acquired along with the green algal plastid and horizontally transferred ones. These findings indicate that euglenophytes’ plastid maturases have experienced a surprisingly dynamic history of their gains, retention, and loss.

Even considering that euglenids stand out among major groups of protists for their abundance of genetic peculiarities (Dobáková et al., 2015; Ebenezer et al., 2019; Karnkowska et al., 2018; Novák Vanclová et al., 2020; Záhonová et al., 2018), our work has shown that their extraordinary capabilities for perplexing researchers are far from being exhausted.

## Abbreviations

GIIM: group II intron maturase
HNHE: HNH endonuclease
IEP: intron-encoded protein
ORF: open reading frame
ptDNA: plastid DNA, plastid genome
RVT: reverse transcriptase/maturase
SSU rDNA: small subunit of the ribosomal DNA

## 5. Acknowledgements

This work was supported by Preludium grant 2018/31/N/NZ8/01840 (National Science Centre, Poland) to KM, Sonata grant 2016/21/D/NZ8/01288 (National Science Centre, Poland) to AK and EMBO Installation Grant to AK. The authors of this manuscript would like to thank Richard E. Triemer and Matthew S. Bennett (Michigan State University, USA) for sharing their unpublished sequencing data from strain *Eutreptiella* sp. CCMP1594.

## 6. Author contributions

AK conceptualized and administrated the project, AK and KM obtained funding, wrote the original draft and prepared figures. KM and ND obtained, curated and analysed the raw data. AK and AP supervised the work. All authors established the methodology to be used (including software), validated the data, and edited, reviewed, and corrected the final manuscript.

## Appendix A –

### supplementary figures

**Figure A.1.**
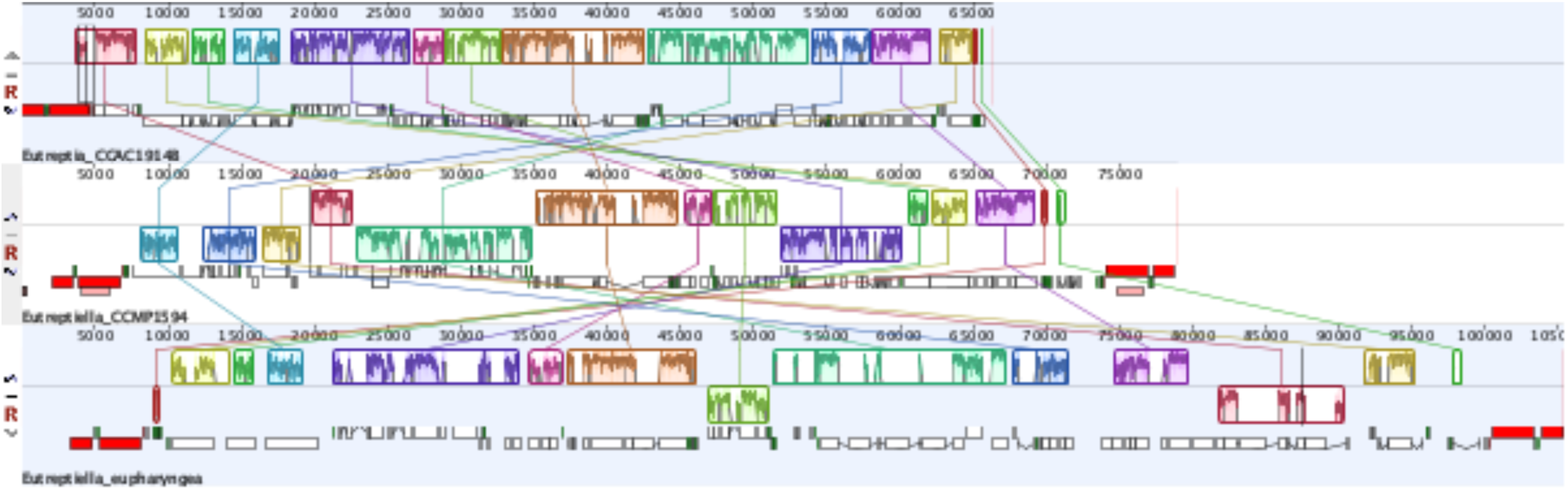
Synteny graph of the newly-sequenced chloroplast genomes of Eutreptiales, generated using Mauve plugin embedded within Geneious software v10.2.6 (Kearse et al., 2012). Note: plastid genome of *Eutreptiella* sp. CCMP389 is not shown, as it is fully syntenic to *Eutreptiella* sp. CCMP1594

**Figure A.2.**
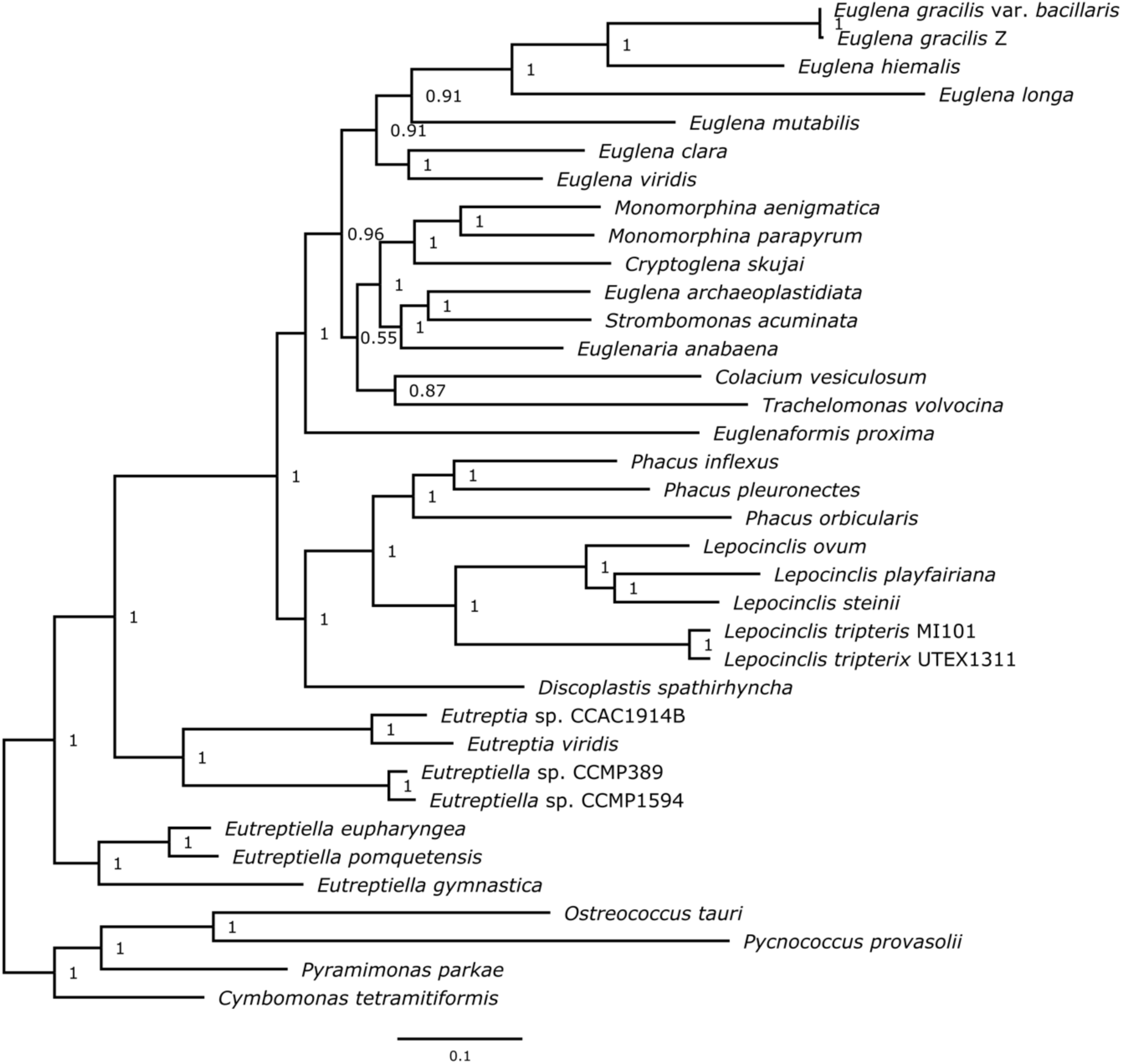
Plastid-based Bayesian phylogenetic tree of Euglenophyta.

**Figure A.3.**
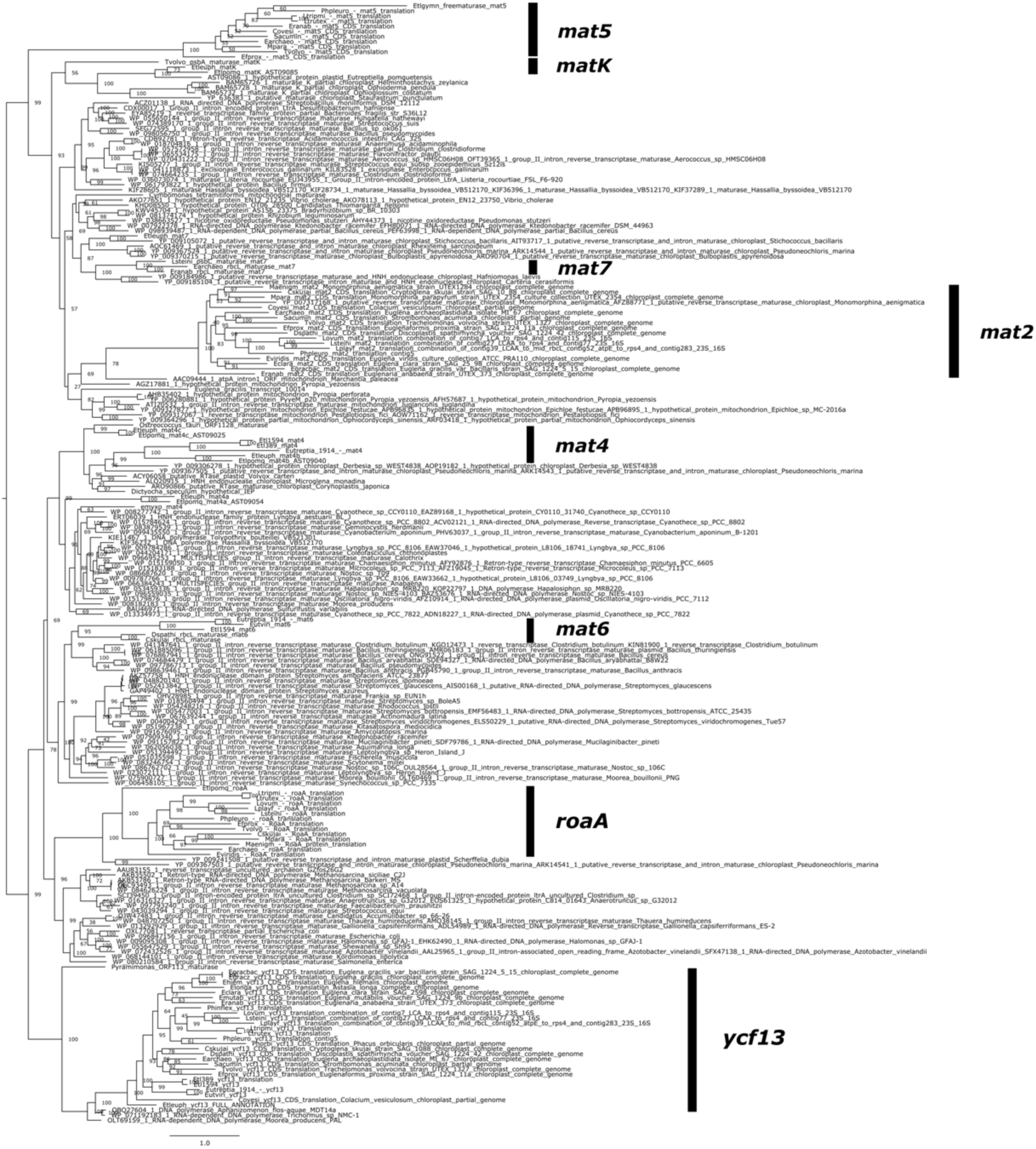
Maximum likelihood phylogenetic tree of euglenid plastid-encoded intron maturases and their known homologs.

**Figure A.4.**
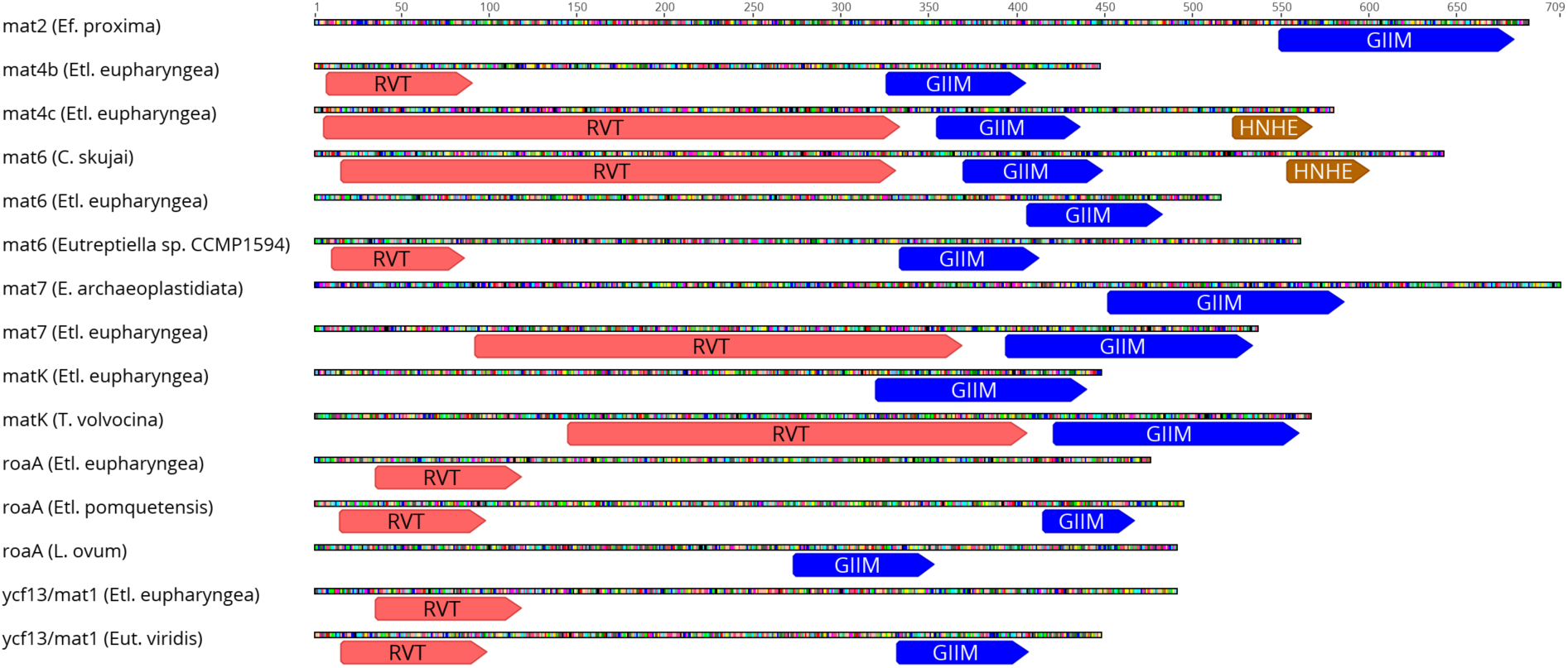
Conserved protein domains in euglenid plastid-encoded maturases, visualized in selected examples. The figure was generated using Geneious software v10.2.6 (Kearse et al., 2012).

## Appendix B – supplementary tables

**Table B.1.** List of published sequences, downloaded from the NCBI GenBank database, used in this study, with their respective accession numbers.

| Accession no. | Name |
| --- | --- |
| AJ294725 | *Astasia longa* complete chloroplast genome |
| AY290861 | *Euglena myxocylindracea* chlorophyll a-binding protein (psbC) gene, partial cds; tRNA-Leu (trnL) gene, complete sequence; photosystem II protein D (psbA), maturase (mat4), and hypothetical chloroplast protein 65 (ycf65) genes, complete cds; tRNA-Arg (trnR) gene, complete sequence; and photosystem I protein M (psaM), hypothetical chloroplast protein 12 (ycf12), and PsbK (psbK) genes, complete cds; chloroplast genes for chloroplast products |
| FJ493498 | *Pycnococcus provasolii* chloroplast, complete genome |
| JN643723 | *Eutreptia viridis* chloroplast, partial genome |
| JN674636 | *Colacium vesiculosum* chloroplast, partial genome |
| JN674637 | *Strombomonas acuminata* chloroplast, partial genome |
| JQ237893 | *Euglena viridis* culture-collection ATCC:PRA110 chloroplast, complete genome |
| JX457480 | *Monomorphina aenigmatica* strain UTEX1284 chloroplast, complete genome |
| KC684276 | *Euglenaformis proxima* strain SAG 1224-11a chloroplast, complete genome |
| KF285527 | *Ostreococcus tauri* isolate RCC1116 chloroplast, complete genome |
| KP410781 | *Cryptoglena skujai* strain SAG 10.88 chloroplast, complete genome |
| KP453743 | *Euglenaria anabaena* strain UTEX 373 chloroplast, complete genome |
| KP455987 | *Monomorphina parapyrum* strain UTEX 2354 culture-collection UTEX:2354 chloroplast, complete genome |
| KP686076 | *Euglena gracilis* var. *bacillaris* strain SAG1224-5/15 chloroplast, complete genome |
| KP686077 | *Trachelomonas volvocina* strain UTEX 1327 chloroplast, complete genome |
| KP939040 | *Euglena archaeoplastidiata* isolate MI 67 chloroplast, complete genome |
| KR921747 | *Phacus orbicularis* chloroplast, partial genome |
| KT223519 | *Euglena mutabilis* voucher SAG 1224-9b chloroplast, complete genome |
| KX013545 | *Cymbomonas tetramitiformis* strain PLY262 chloroplast, complete genome |
| KX013546 | *Pyramimonas parkeae* strain NIES254 chloroplast, complete genome |
| KY706202 | *Eutreptiella pomquetensis* plastid, partial genome |
| MF622086 | *Euglena hiemalis* strain CCAP1224.35 chloroplast, complete genome |
| MF630936 | *Euglena clara* strain SAG 25.98 chloroplast, complete genome |
| MH898667 | *Phacus inflexus* voucher ACOI 1336 chloroplast, complete genome |
| MH898668 | Lepocinclis tripteris voucher MI 101 chloroplast, complete genome |
| MH898669 | Lepocinclis tripteris voucher UTEX 1331 chloroplast, complete genome |
| MH898670 | Discoplastis spathirhyncha voucher SAG 1224-42 chloroplast, complete genome |
| MH898671 | Lepocinclis playfairiana voucher MI 102 chloroplast, complete genome |
| MH898672 | Lepocinclis steinii voucher UTEX 523 chloroplast, complete genome |
| MH898673 | Phacus pleuronectes voucher SAG 1261-3b chloroplast, complete genome |
| MH898674 | Lepocinclis ovum voucher SAG 1244-8 chloroplast, complete genome |
| NC_01775 | Eutreptiella gymnastica plastid, complete genome |
| X70810 | Euglena gracilis chloroplast complete genome |

